# Precise and stable edge orientation signaling by human first-order tactile neurons

**DOI:** 10.1101/2022.06.01.494420

**Authors:** Vaishnavi Sukumar, Roland S. Johansson, J. Andrew Pruszynski

**Author notes:** **Corresponding Author** J. Andrew Pruszynski, Robarts Research Institute, Room 1232H, Western University, London, Ontario, Canada, N6A 5B7, E, T: @andpru.

## Abstract

Fast-adapting type 1 (FA-1) and slow-adapting type 1 (SA-1) first-order neurons in the human tactile system have distal axons that branch in the skin and form many transduction sites, yielding receptive fields with many highly sensitive zones or ‘subfields’. We previously demonstrated that this arrangement allows FA-1 and SA-1 neurons to signal the geometric features of touched objects, specifically the orientation of raised edges scanned with the fingertips. Here we show that such signaling operates for fine edge orientation differences (5-20°) and is stable across a broad range of scanning speeds (15-180 mm/s); that is, under conditions relevant for real-world hand use. We found that both FA-1 and SA-1 neurons weakly signal fine edge orientation differences via the intensity of their spiking responses and only when considering a single scanning speed. Both neuron types showed much stronger edge orientation signaling in the sequential structure of the evoked spike trains and FA-1 neurons performed better than SA-1 neurons. Represented in the spatial domain, the sequential structure was strikingly invariant across scanning speeds, especially those naturally used in tactile spatial discrimination tasks. This speed invariance suggests that neurons’ responses are structured via sequential stimulation of their subfields and thus links this capacity to their terminal organization in the skin. Indeed, the spatial precision of elicited action potentials rationally matched spatial acuity of subfield arrangements, which typically corresponds to the dimension of individual fingertip ridges.

**Significance Statement:** The distal axons of human first-order tactile neurons branch and innervate many mechanosensitive end organs in the skin. For those neurons terminating in end organs associated with fingerprint ridges (Meissner and Merkel), this branching results in cutaneous receptive fields with multiple subfields spread across several ridges. Consequently, when a fingertip scans the surface of an object, the spatial coincidence between a neuron’s subfields and the tactile stimulus defines the sequential structure of the evoked spike train (i.e., the presence of action potential bursts and the gaps between them). Here we show that, for surfaces composed of oriented edges, this sequential structure signals information about edge orientation differences at the limit of what people can feel and that the spatial precision of the structuring is maintained across a broad range of speeds relevant for real-world hand use. We submit that, to be of human relevance, models of higher order tactile processing must consider the impact of multifocal receptive fields in the periphery. For example, the speed invariance of tactile fine-form/texture perception may arise simply because the same subsets of peripheral subfields in the population of first-order tactile neurons are stimulated together regardless of speed.

## Introduction

The distal axons of fast-adapting type 1 (FA-1) and slow-adapting type 1 (SA-1) neurons innervating the glabrous skin of the primate hand branch extensively in the skin, causing each individual neuron to innervate many spatially segregated low-threshold mechanoreceptive end organs (Cauna, 1956, 1959; Nolano et al., 2003; Paré et al., 2002; Vallbo & Johansson, 1984). This arrangement results in spatially complex receptive fields with many highly sensitive zones (or ‘subfields’) distributed within a circular or elliptical area typically covering five to ten fingerprint ridges (Jarocka et al., 2021; Johansson, 1978; Phillips et al., 1992; Pruszynski & Johansson, 2014). We have proposed that this arrangement constitutes a peripheral neural mechanism for geometric feature extraction (Hay & Pruszynski, 2020; Jarocka et al., 2021; Pruszynski et al., 2018; Pruszynski & Johansson, 2014; Zhao et al., 2018). Consistent with this idea, we previously demonstrated that both FA-1 and SA-1 neurons signal information about the orientation of edges moving across a neuron’s receptive field via changes in both their spiking intensity and the sequential structure of their spiking response (Pruszynski & Johansson, 2014). The intensity of a neuron’s response can be modulated by the degree of spatial coincidence between the neuron’s subfields and local skin deformation caused by the edge. The sequential structure of a neuron’s response (i.e., the gaps between bursts of spikes) can carry information about edge orientation because the orientation affects the sequence and timing of stimulated subfields as the edge passes over the neuron’s receptive field.

Here we examine this peripheral signaling mechanism with respect to the ability of individual FA-1 and SA-1 neurons to signal fine differences in the orientation of edges (5 – 20°) moving across the fingertip and how stimulation speed affects this ability. In our previous study (Pruszynski & Johansson, 2014) we used edges that had large orientation differences (≥22.5°) compared to edge orientation discrimination thresholds found during object manipulation (∼3°, Pruszynski et al., 2018) or psychophysical tasks (∼10-25°, Bensmaia et al., 2008; Lechelt, 1992; Olczak et al., 2018; Peters et al., 2015) and we focused mainly on one movement speed (30 mm/s). However, during natural contacts with objects where the use of tactile information is critical, a wide range of speeds can occur(Callier et al., 2015; Cole & Abbs, 1988; Johansson & Westling, 1984; Olczak et al., 2018; Smith, Chapman, et al., 2002; Smith, Gosselin, et al., 2002; Vega-Bermudez et al., 1991). This natural variation motivated us to analyze the capacity of individual neurons to signal edge orientation at different fixed speeds but also their ability to signal edge orientation across scanning speeds (i.e., in a speed-invariant manner). Examining speed invariance also allowed us to test the hypothesis that the subfield arrangement of FA-1 and SA-1 neurons gives rise to their ability to signal fine edge orientation via changes in the sequential structure of their spiking responses. Because the layout of a neuron’s subfield arrangement is fairly stable over a range of natural scanning speeds (Jarocka et al., 2021), the hypothesis predicts speed invariant signaling of edge orientation when spike trains are represented in the spatial domain, where the action potentials are represented with respect to the position of the stimulus on the skin at the time of their occurrence.

We recorded action potentials in the distal axon of single FA-1 and SA-1 neurons innervating human fingertips while tangentially scanning their receptive field with finely oriented edges (±5° and ±10° relative to the scanning direction) at speeds spanning more than one order of magnitude (15 - 180 mm/s). We report that FA-1 and SA-1 neurons can signal fine edge orientation differences via changes in spiking intensity (peak and mean firing rate) but that they do so much more reliably by changes in the sequential structure of the evoked spike trains. In the latter case, FA-1 neurons perform better than SA-1 neurons, but both show best edge orientation discrimination capacity for scanning speeds that humans naturally use when moving their hand to discriminate fine spatial features. Furthermore, when spike trains are represented in the spatial domain, their orientation-dependent sequential structuring is preserved across a wide range of scanning speeds, meaning that FA-1 and SA-1 neurons can provide speed invariant information about edge orientation. Taken together, these findings further the idea that the dendritic-like branching of first-order tactile neurons’ distal axons contributes to early processing of fine geometric features in the tactile sensory ascending pathways.

## Methods

### Study participants and data sample

Human participants (5 male, 8 female) provided written informed consent in accordance with the Declaration of Helsinki. The Umeå University ethics committee approved the study. The general experimental methodology, procedure and apparatus have been described previously (Pruszynski & Johansson, 2014).

We recorded action potentials from single first-order tactile neurons terminating in the glabrous skin of the tips of the index, long or middle fingertips, using tungsten electrodes (Vallbo & Hagbarth, 1968). The electrodes were inserted into the median nerve at the level of the upper arm or wrist. The present study focuses on fast adapting type-1 (FA-1) or slow adapting type-1 (SA-1) neurons, classified according to previously described criteria (Vallbo & Johansson, 1984). Of the 53 neurons recorded, 30 were FA-1 and 23 were SA-1.

### Stimuli

Each neuron was stimulated by lines embossed on a flat surface that moved tangentially across the receptive field along the proximal distal axis of the finger (**Figure 1A**). The stimulation surface was a photo-etched nylon polymer (EF 70 GB, Toyobo Company, Japan) wrapped around a rotating drum (diameter = 59 mm).

**Figure 1.**
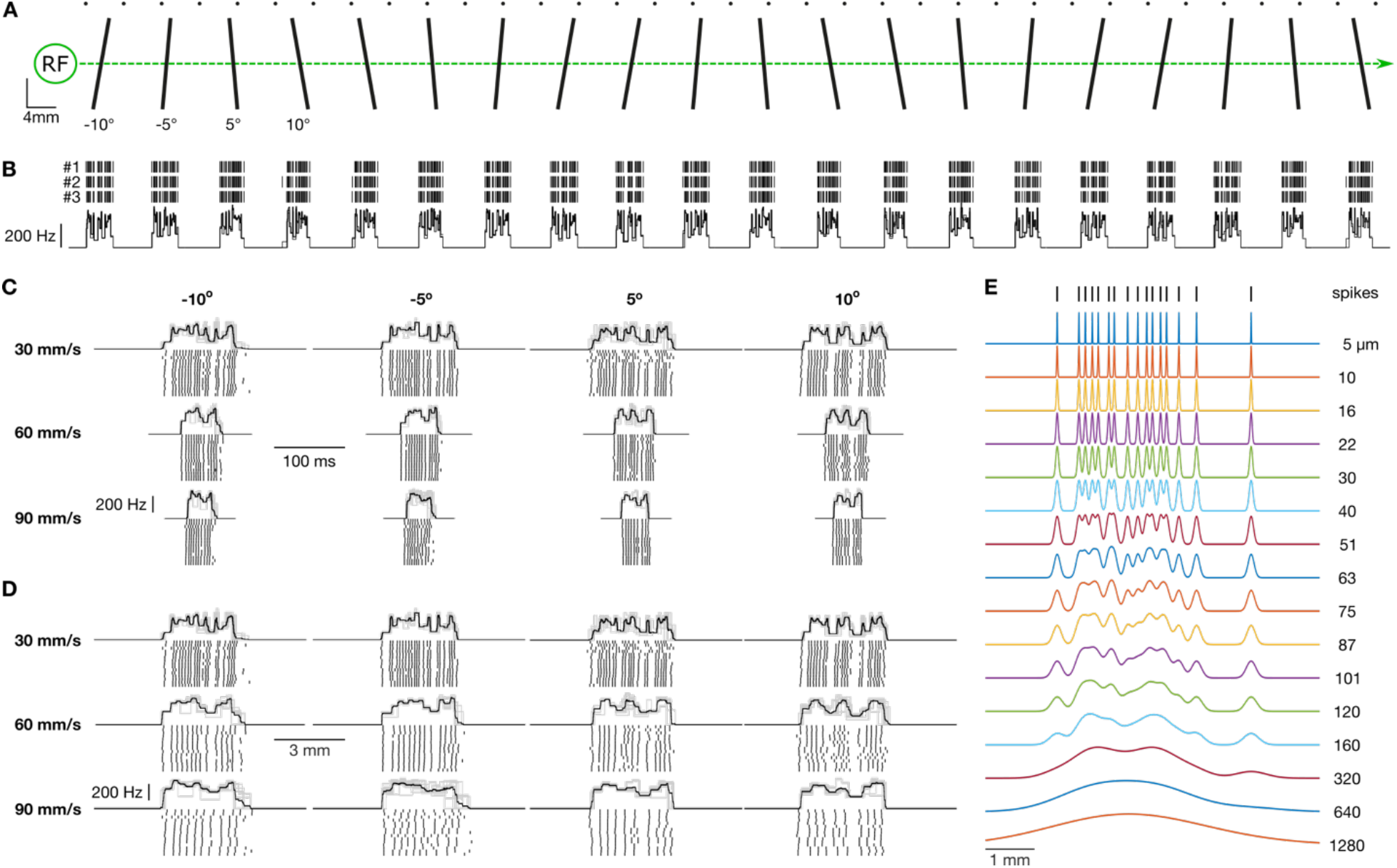
**(A)** Schematic of the stimulating surface, which includes five repetitions of four oriented line stimuli. The small dots above the line stimuli are used during the experiment to align the stimulating surface with an isolated neuron’s receptive field (RF). The layout of the line stimuli is such that an isolated RF is only stimulated by a single line stimulus at any given moment in time. **(B)** Tick marks represent action potentials from an exemplar neuron plotted relative to the position of the stimulating surface. All three rotations (#1, #2, #3) are shown. The superimposed traces below represent the firing rate profile, defined as the reciprocal of the interval between subsequent action potentials, for each of the three rotations. This constitutes the raw data for this study. **(C)** Response from an exemplar neuron for the four line orientations and three stimulating speeds. The black trace represents the mean response and the light gray traces show responses for the 15 individual trials. Tick marks below represent action potentials for each individual trial. Scale bars show time (100 ms). **(D)** Same data as in C but represented in spatial coordinates. Scale bars show distance traversed by the drum (2 mm). **(E)** Each trace shows an exemplar spike train smoothed by Gaussian kernels of various widths (noted is the standard deviation the kernel).

A custom-built robotic device controlled the rotation speed of the drum (see Pruszynski & Johansson, 2014). To stabilize the fingertips, the nails were glued to plastic holders firmly attached to a table that also supported the right arm and the robot and the contact force perpendicular to the skin was servo-controlled to ∼0.4 N. The instantaneous rotational position of the drum was monitored via an optical shaft encoder (AC36, Hengstler GmbH), which provided position resolution of 3 µm in the scanning direction.

Raised lines oriented -10°, -5°, 5° and 10° relative to the axis perpendicular to the direction of motion were presented five times on the stimulus pattern (**Figure 1A**). The lines, extending across the width of the stimulus surface (12 mm), were 0.5 mm high and their widths were 0.5 mm at the top and 0.8 mm at the base. All lines were spaced at least 8 mm apart to minimize interactions between neighboring lines on a neuron’s response.

A neuron’s receptive field was scanned at up to twelve different tangential speeds: 2.5, 5, 10, 15, 20, 30, 45, 60, 90, 120, 180 mm/s in random order. For each neuron and speed, the drum rotated three times, resulting in 15 trials for each line stimulus in speed conditions recorded (5 presentations of stimuli per drum rotation x 3 drum rotations = 15 trials).

## Data processing and analysis

### Sampling

The nerve signal, the instantaneous position of the stimulus surface and the contact force were digitally sampled at 19.2 kHz, 2.4 and 0.6 kHz respectively (SC/ZOOM, Department of Integrative Medical Biology, Umeå University).

To quantify the intensity of a neuron’s response we used peak and mean firing rate. For each spike train evoked by an edge passing over the receptive field, we calculated peak firing rate as the reciprocal of the shortest interspike interval observed in the evoked response. Mean firing rate was calculated as the number of spikes evoked within an 8 mm response window (the minimum distance between adjacent elements on the drum) divided by the duration of the stimulus traversing this window. Furthermore, for each spike train we generated the firing rate profile by computing the instantaneous frequency of the action potentials defined as the inverse of the interval between consecutive impulses for the duration of the interval. This calculated rate profile was then treated as a continuously recorded signal.

### Orientation signaling as a function of scanning speed

To analyze how well a neuron signaled edge orientation at each scanning speed based on the above response measures, we used methods analogous to those previously described (Pruszynski & Johansson, 2014).

For the intensity measures, we calculated how frequently the peak or mean firing rate evoked by each presentation of a particular line orientation was closer to the average response of the remaining (14) trials with the same orientation, as opposed to the average response of 14 trials from the other three edge orientations (chance = 0.25). By including only 14 of the 15 trials from other stimuli, we avoided a bias caused by a better estimate of the average response to the other stimuli.

For a neuron’s firing rate profiles, we calculated the probability of correctly classifying an observed profile to the line stimulus that evoked it. To that end, we pairwise cross-correlated all firing rate profiles obtained at the speed in question. For each profile, we then calculated an average of the correlations between the remaining (14) trials with the same line orientation and the average for 14 trials from each of the other three line-orientations. Finally, we calculated how often the highest average correlation resulted from stimuli with the same line orientation as opposed to the other orientations (chance = 0.25). We again only considered 14 other trials to avoid classification bias.

### Speed-invariant orientation signaling

To investigate a neuron’s capacity to signal edge orientation in a speed invariant manner, we applied the classification approach described above for within-speed signaling of other speed conditions. For the intensity measures, we quantified how frequently the peak or mean firing rate evoked by each presentation of a particular line orientation at one speed was closer to the average response of the trials with the same line orientation for another speed (i.e., correctly classified) as opposed to trials with the other line orientations (i.e., incorrectly classified). For firing rate profiles, we quantified how often a neuron’s firing rate profile for a given orientation and speed was best correlated with profiles evoked by stimuli with the same orientation but at different speeds (i.e., correctly classified) as opposed to profiles evoked with the other line orientations also at different speeds (i.e., incorrectly classified). These analyses were performed with firing rate profiles conventionally represented in the temporal domain but also represented in the spatial domain where the positions of the underlying action potential are referenced to the position of the stimulus surface relative to the skin.

### Precision of spiking responses

The hypothesis that subfield arrangement structures sequentially a neuron’s spiking activity when edges pass over their receptive fields motivated us to estimate the spatial precision of generated action potentials with reference to neurons’ signaling of fine edge orientation differences. For this purpose, we used a correlation-based measure of similarity between pairs of individual spike trains after imposing various amounts of noise on the position of the recorded action potentials (Fellous et al., 2004; Schreiber et al., 2003). Thus, we convolved each spike train with Gaussians of different kernels with roughly logarithmically spaced standard deviations in the range of 5-1280 µm, which corresponds to increasingly blurring a neuron’s receptive field sensitivity topography (Jarocka et al., 2021). For each neuron and kernel width, we then pairwise correlated all smoothed spike trains and determined the spatial precision of action potentials by calculating which kernel width yielded the largest average difference in correlations with the same orientations at other speeds and with different orientations at different speeds (termed “best kernel”). This analysis thus only considered correlations between speed conditions since our hypothesis predicts speed invariant edge orientation signaling when spike trains are represented in the spatial domain.

For spike trains convolved with the best kernel averaged across speeds for the FA-1 and SA-1 neurons, respectively, we also quantified for each neuron type the proportion of correctly classified orientations across scanning speeds analogous to the analysis of speed-invariant orientation signaling with firing rate profiles (chance performance = 0.25). Finally, based on pairwise comparisons between the available stimuli, we quantified the proportion of correctly classified orientations as a function of angular difference (chance = 0.5). Given our data set, the available angular differences were 5° (between -5° and -10° and between +5° and +10°), 10° (between -5° and +5°), 15° (between -10° and +5° and between -5° and +10°) and 20° (between -10° and +10°).

### Statistics

We carried out all statistical analysis in R Studio (V1.2.5). Details of statistical analysis are provided in the text including the degrees of freedom, the test statistic, the p-value, and the effect size (partial eta squared: ηp^2^). In all cases, kernel width data was log-transformed, and discrimination probability data was arcsine square root transformed where needed. For ANOVAs evaluating discrimination probability, we subtracted chance from the data such that a significant intercept in the ANOVA model meant that overall performance was above chance. These analyses included scanning speed (and sometimes orientation difference) as a within-group (repeated measures) factor and neuron type as a between-group factor.

## Results

We examined the capacity of individual FA-1 and SA-1 neurons innervating human fingertips to signal fine orientation differences of edges moving across their receptive field as a function of scanning speed as well as their ability to signal fine orientation differences in a speed-invariant manner. We did so with respect to different aspects of their spiking responses. These included traditional intensity measures (i.e., peak and mean firing rate) as well as spatiotemporal measures based on the sequential structure of the spike trains. For the sequential structure, our emphasis was on its representation in the spatial domain where each action potential is referenced to the position of the edge stimulus on the skin when it occurred as this relates to the spatial structuring of the neuron’s receptive field in the form of subfields.

We recorded action potentials from 30 FA-1 and 23 SA-1 isolated neurons when stimulated by repeatedly sweeping raised lines across their receptive field (**Figure 1A**). The lines were oriented ±5° and ±10° relative to their direction of motion. All neurons were stimulated at eight speeds from 15 to 180 mm/s. Each edge orientation was presented 15 times per speed. The contact force between the stimuli and the finger was maintained at ∼0.4 N, which is typical for haptic exploration with such stimuli (Olczak et al., 2018). **Figure 1B** shows the response of an exemplar neuron to the edges as a function speeds. The scanning speed affected neurons’ responses as expected. That is, mean firing rate (**Figure 2A**) and peak firing rate (**Figure 2B**) increased with speed and the number of spikes decreased with speed (**Figure 2C**); in general, FA-1 neurons responded with higher rates than SA-1 neurons.

**Figure 2.**
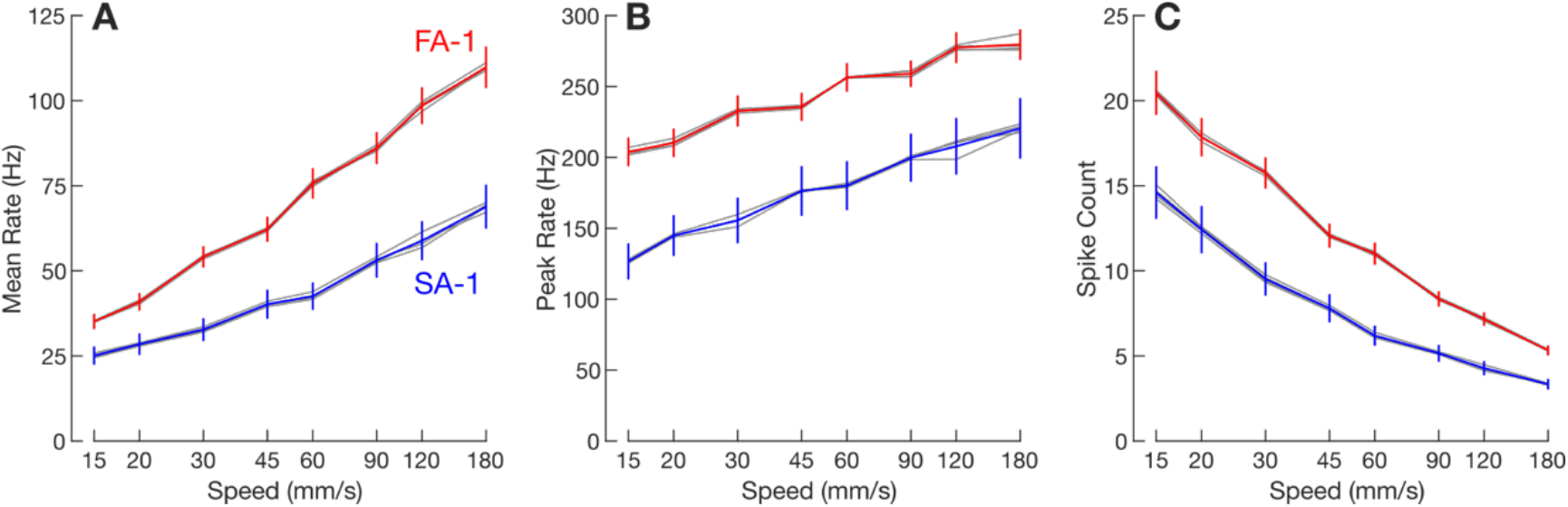
Response intensity as a function of all stimulating speeds used in this study. **(A)** Mean firing rate averaged across FA-1 neurons (red line) and SA-1 neurons (blue line), respectively. The error bars represent the standard error of the mean across neurons (FA-1: n = 30; SA-1: n = 23). The thin gray lines represent mean responses for each of the four line stimuli separately. **(B**,**C)** Same format but for peak firing rate and spike count, respectively.

### Orientation signaling as a function of scanning speed

For each scanning speed we calculated how well an ideal observer could correctly classify an edge orientation based on a given neuron’s peak and mean firing rate (see Methods). Discrimination accuracy based on both of these measures was poor but above chance across the speeds (**Figure 3A,B**). That is, separate two-way ANOVAs with speed as a within-group (repeated measures) factor and neuron type as a between-group factor, applied to the proportion of correctly classified edge orientations referenced to chance performance (25%), revealed a significant intercept for both peak (F_1,51_ = 556.9, p < 0.001, ηp^2^ = 0.92) and mean (F_1,51_ = 196.3, p < 0.001, ηp^2^ = 0.79) firing rate.

**Figure 3.**
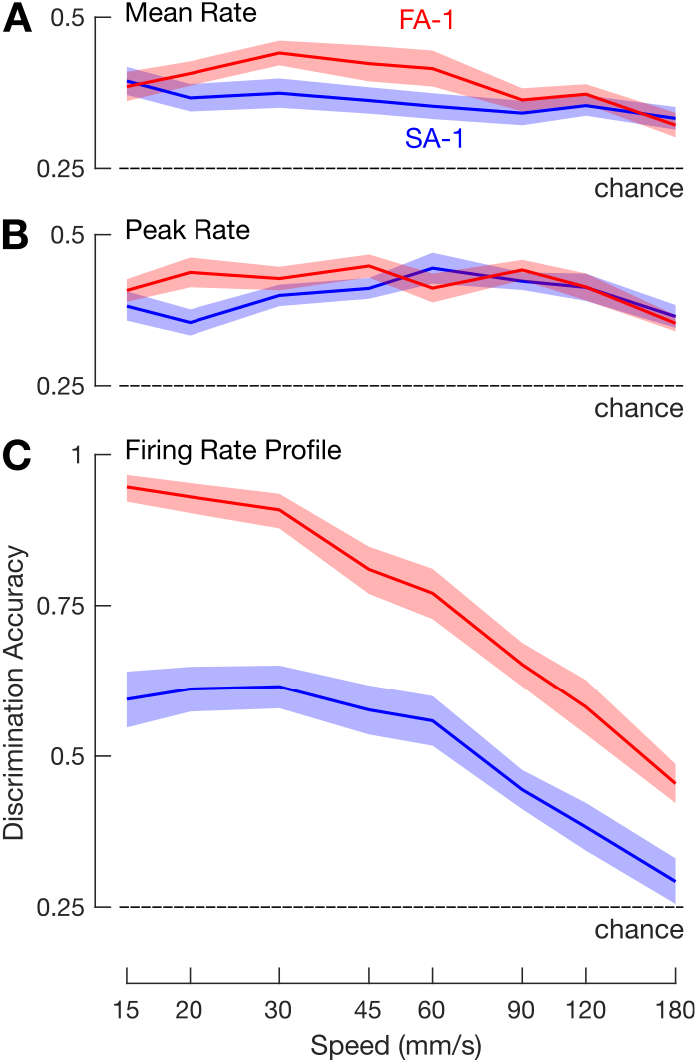
Average (mean) within-speed discrimination accuracy for FA-1 neurons (red lines) and SA-1 neurons (blue lines) as a function of simulation speed based on neuron’s mean firing rate (**A**), peak firing rate (**B**) and firing rate profile (**C**). **(A** – **C**) Shaded areas illustrate the standard error of the mean and dashed line chance discrimination performance (25%).

For peak firing rate, mean discrimination accuracy averaged across neurons and scanning speeds was 42% (range as a function of speed: 35-44%) for FA-1 neurons and 39% (range: 35-44%) for SA-1 neurons. We found no significant effect of neuron type on discrimination accuracy (F_1,51_ = 1.7, p = 0.20, ηp^2^ = 0.03) but observed a reliable effect of speed (F_7,357_ = 3.4, p < 0.01, ηp^2^ = 0.06), indicating a tendency for worse discrimination with increasing speed. For mean firing rate, mean discrimination accuracy averaged across the scanning speeds was 39% (range as a function of speed: 32-44%) for FA-1 neurons and 36% (range: 33-39%) for SA-1 neurons. Again, we found no significant effect of neuron type on discrimination accuracy (F_1,51_ = 2.6, p = 0.11, ηp^2^ = 0.05) but an effect of speed (F_7,357_ = 4.1, p < 0.001, ηp^2^ = 0.07) indicating a tendency for worse discrimination with increasing speed. For both peak and mean firing rate, the effect of speed might relate to the decreasing number of action potentials generated per edge stimulus at higher speeds.

Given that edge orientation signaling can occur via changes in the sequential structure of a neuron’s spiking response (Pruszynski & Johansson, 2014), we calculated how well an ideal observer could correctly classify edge orientation based on a neuron’s firing rate profile as a function of scanning speed (see Methods). Discrimination accuracy based on firing rate profile was substantially higher than that based on intensity measures (**Figure 3C**). Averaged across the speeds, mean discrimination accuracy was 78% for FA-1 neurons (range as a function of speed: 45-95%) and 51% for SA-1 neurons (range: 29-62%). A two-way ANOVA applied to the proportion of correctly classified edge orientations revealed a significant intercept referenced to chance (25%) performance (F_1,51_ = 309.5, p < 0.001, ηp^2^ = 0.86). Discrimination accuracy increased with decreasing speed (F_7,357_ = 71.1, p < 0.001, ηp^2^ = 0.58) and was, on average, better for FA-1 neurons than for SA-1 neurons (F_1,51_ = 33.2, p < 0.001, ηp^2^ = 0.40). The superiority of FA-1 neurons was at low to moderate speeds while the discrimination accuracy of both neuron types approached chance performance at the highest speeds, giving rise to a significant interaction between speed and neuron type (F_7,357_ = 6.5, p < 0.001, ηp^2^ = 0.11).

These results show that the sequential structure of individual first-order tactile neuron responses contains considerable information about fine edge orientation differences. However, it is not obvious that neuron subfield arrangements are a crucial factor in the structuring of responses in this regard. That is, first-order tactile neurons can be incredibly sensitive to skin distortions and produce remarkably repeatable spiking responses with temporal precision to the millisecond level, so it may be argued that edge orientation discrimination can be based on any difference in mechanical events produced by non-identical stimuli (Suresh et al., 2016).

### Speed-invariant orientation signaling

The following section deals with the ability of individual first-order tactile neurons to precisely signal information about edge orientation in a speed-invariant manner. We are motivated to investigate speed-invariance not only because speed often varies when touching and manipulating objects but also because it provides one means of testing whether the subfield arrangement of FA-1 and SA-1 neurons is an important factor for their signaling of fine differences in edge orientation.

Based on a neuron’s response to a line stimulus recorded at a given speed, we calculated how well an ideal observer could identify the correct edge orientation based on the neuron’s responses at the other speeds (see Methods). As expected, intensity measures (i.e., peak and mean firing rates), which were strongly affected by the scanning speed (**Figure 2**), failed to provide any reliable information about edge orientation at other scanning speeds (intercept of two-way ANOVA referenced to chance performance; peak firing rate: F_1,51_ = 0.09, p = 0.77, ηp^2^ = 0.002; mean firing rate: F_1,51_ = 0.07, p = 0.72, ηp^2^ = 0.001; **Figure 4A,B**). Also as expected, given that a neuron’s spike train is compressed and dilated in time as a function of speed (**Figure 1B**), firing rate profiles represented in the temporal domain did not retain reliable information about edge-orientation at other stimulation speeds (F_1,51_ = 0.59, p = 0.44, ηp^2^ = 0.01; **Figure 4C**).

**Figure 4.**
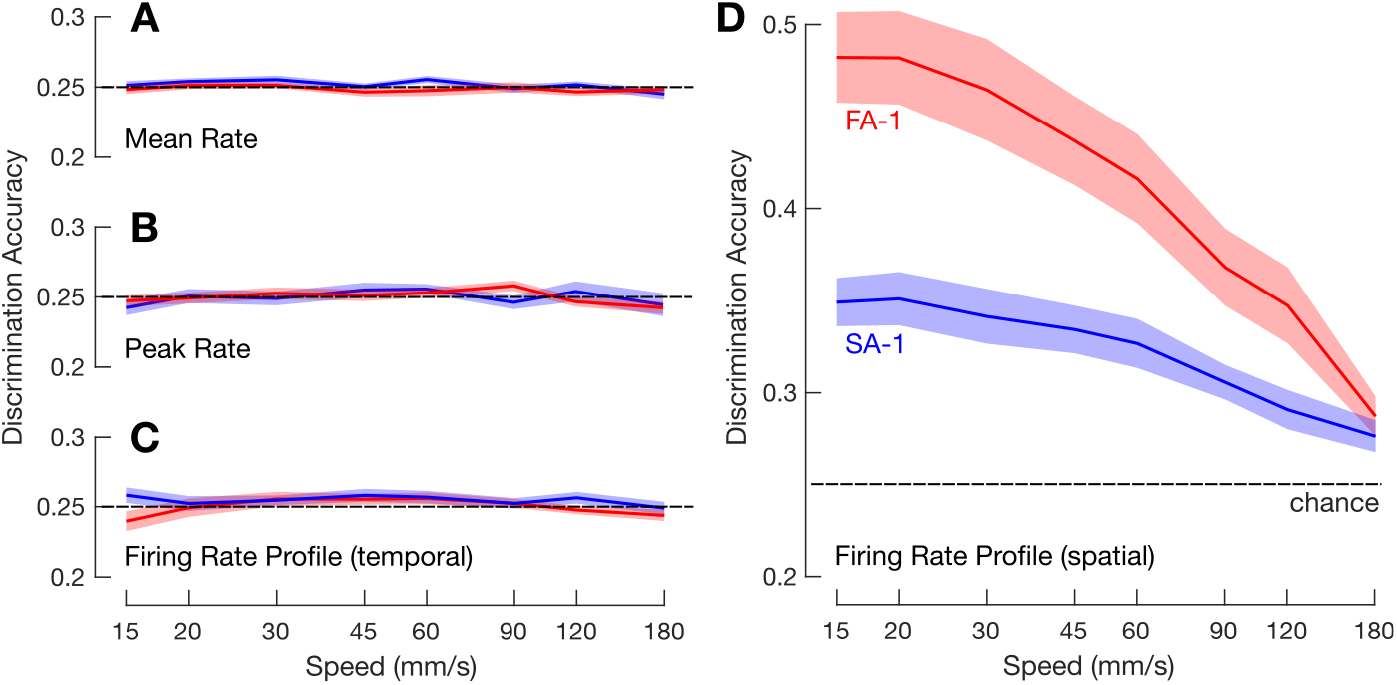
Mean across-speed discrimination accuracy for FA-1 neurons (red lines) and SA-1 neurons (blue lines) as a function of core speed based on neuron’s mean firing rate (**A**), peak firing rate (**B**), firing rate profile represented in the temporal domain (**C**), and in the spatial domain (**D**). (**A** – **C**) Shaded areas illustrate the standard error around the mean (FA-1: n = 30; SA-1: n = 23) and dashed horizontal line chance discrimination performance (25%).

In contrast, firing rate profiles contained substantial information about edge orientation at other speeds if represented in the spatial domain and therefore not compressed or dilated with speed changes (i.e., when the position of each action potential is referenced to the position of the edge relative to the skin; **Figure 4D**). Averaged across the speeds, mean discrimination accuracy was 41% (range as a function of speed: 29-48%) among FA-1 neurons and 32% (range: 28-35%) among SA-1 neurons. A two-way ANOVA referenced to chance performance (25%), with speed as a within-group factor and neuron type as a between-group factor, revealed a significant intercept, (F_1,51_ = 119.2, p < 0.001, ηp^2^ = 0.70). Across-speed discrimination accuracy was reliably better for FA-1 neurons than for SA-1 neurons (F_1,51_ = 14.2, p < 0.001, ηp^2^ = 0.22). Discrimination accuracy also varied with speed (F_7,357_ = 41.9, p < 0.001, ηp^2^ = 0.45), with a decrease in performance for higher speeds presumably related to the relative decrease in the overall number of action potentials per stimulus. A significant interaction between speed and neuron type (F_7,357_ = 6.5, p < 0.01, ηp^2^ = 0.11) reflected a higher increase in discrimination accuracy with decreasing speed for the FA-1 than for SA-1 neurons.

### Spatial precision of spike responses

The analysis above indicates that the spatial structuring of FA-1 and SA-1 responses is fairly stable over scanning speeds, which supports our hypothesis that neurons’ subfield arrangements gives rise to their ability to signal fine edge orientation differences. However, it does not establish whether the spatial precision of the generated action potentials aligns with the spatial acuity of the subfield arrangement of FA-1 and SA-1 neurons (Jarocka et al., 2021).

We assessed the spatial precision of the spiking responses by applying varying amounts of noise to the position of the recorded action potentials and quantifying how the noise affected the neuron’s across-speeds signaling of edge orientation. In practice, each individual spike train was convolved with Gaussian kernels of various widths (5-1280 µm) that increasingly attenuated the spatial structure of the response (**Figure 5A**), analogous to blurring the spatial acuity of the neuron’s subfield arrangement. For each kernel width, we then correlated each of the convolved spike trains obtained for a given edge orientation with (1) the spike trains obtained with the same orientation at all other speeds and (2) the spike trains obtained with the other orientations at all other speeds.

**Figure 5.**
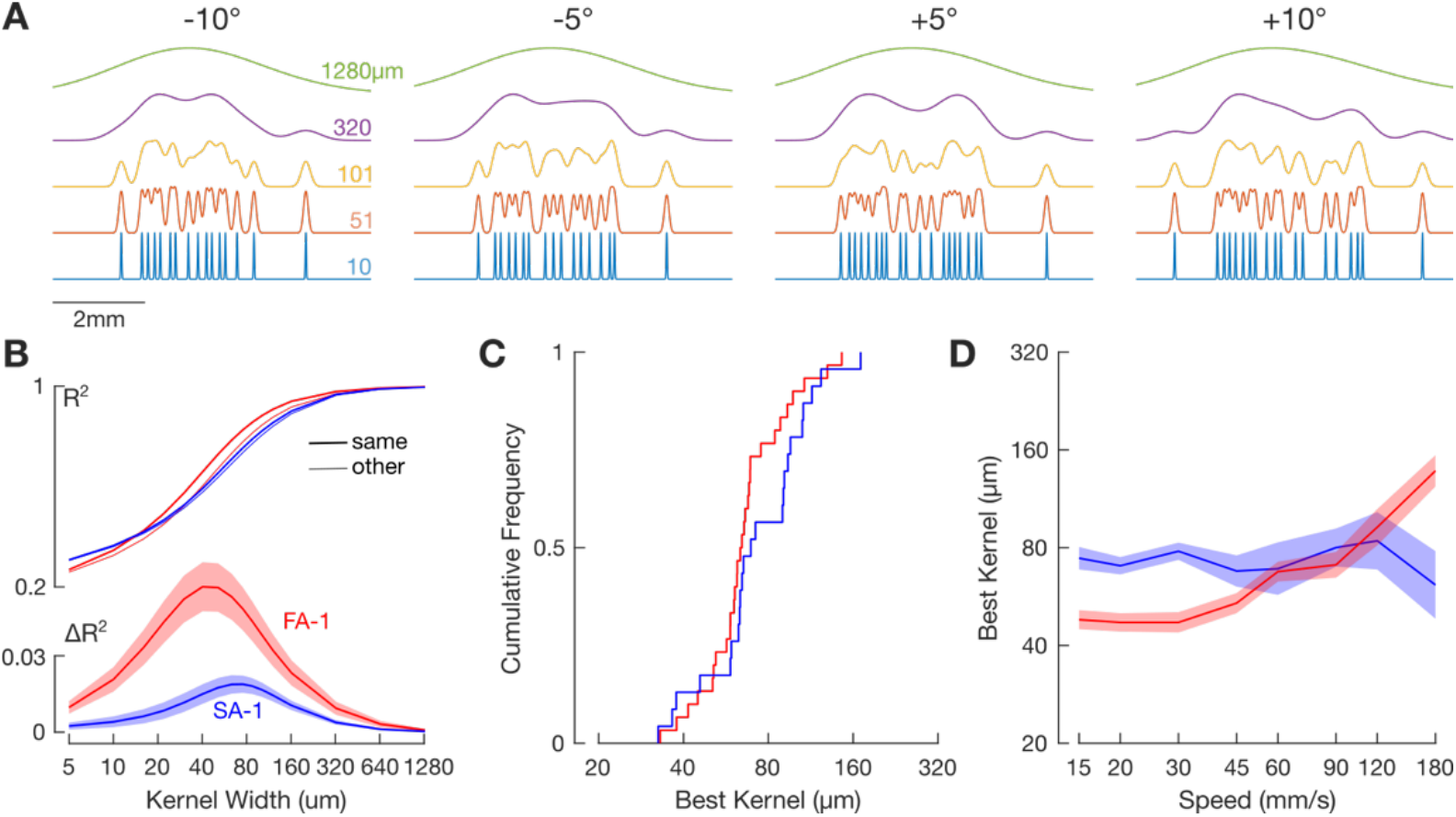
**(A)** Illustration of the kernel convolution procedure for an exemplar neuron for one speed and all four line stimuli. Colored traces within each panel show the spike train smoothed by a Gaussian kernel of the indicated standard deviation. Note how increasing the kernel width progressively blurs the representation of where individual action potentials occur. **(B)** Top: Mean correlation (expressed as R^2^) between neuron responses to the same edge orientations (thick line) and different edge orientations (thin line) as a function of kernel width for the 30 mm/s scanning speed. Bottom: Average correlation difference between stimuli with the same and with different edge orientations as a function of kernel width for the 30 mm/s scanning speed. Red and blue lines represent FA-1 and SA-1 neurons, respectively. Shaded areas illustrate the standard error around the mean. **(C)** Cumulative frequency distribution of the best kernel of FA-1 (red) and SA-1 (blue) neurons where each neuron is represented by the mean value of the best kernel calculated across speeds. **(D)** Mean best kernel as a function of scanning speed for FA-1 (red line) and SA-1 neurons (blue line). Shaded areas illustrate the standard error around the mean (FA-1: n = 30; SA-1: n = 23).

In either case, similarly low correlations arose with the narrowest kernels because the spike jitter between repetitions of the same edge-speed combination tended to be greater than the kernel width. With increasing kernel width, the correlation gradually increased indicating that the spike trains became more similar; the correlation approached one with the widest kernels because all stimulus-dependent structuring of the spike trains was then practically blurred (**Figure 5B**, upper panel). At intermediate kernel widths, however, the correlations between the convolved spike trains originating from different edge orientations were smaller than corresponding correlations between spike trains originating from the same orientations. At these kernel widths the spike trains from different edge orientations were more different with respect to features sensitive to edge orientation differences. Hence, the kernel width at which the difference between these two sets of correlations is greatest provides an estimate of the spatial precision of action potentials critical for edge orientation signaling across speeds.

For each neuron and kernel-speed combination we averaged the correlations involving trials with the same and different edge orientations and then calculated the difference between these averages. This difference was unimodal and followed an inverted-U profile as a function of kernel width, approaching zero when kernels were very narrow and wide (**Figure 5B**, lower panel). The maximum value therefore provided a distinct estimate of a neuron’s spatial precision as a function of speed.

The distribution of best kernels averaged across scanning speeds was narrow with a mean value of 66 µm for FA-1 neurons and 73 µm for SA-1 neurons (**Figure 5C**). A two-way ANOVA showed a main effect of scanning speed (F_7,357_ = 8.38, p < 0.001, ηp^2^ = 0.141) but not of neuron type (F_1,51_ = 0.98, p = 0.32, ηp^2^ = 0.019). There was also a significant interaction between the effects of these factors (F_7,357_ = 8.4, p < 0.001, ηp^2^ = 0.142) as SA-1 kernels were less sensitive to scanning speed (**Figure 5D**). Notably, the estimates of spatial precision based on fine edge orientation discrimination are very similar to the estimates of spatial acuity of the subfield arrangements for FA-1 and SA-1 neurons (see Jarocka et al., 2021 and Discussion).

### Discrimination accuracy based on convolved spike trains

Here we examined across-speed discrimination accuracy based on convolved spike trains. We calculated discrimination accuracy across the speeds after convolving the spike trains with the mean of the best kernel across FA-1 and SA-1 neurons averaged across the speeds (66 and 73 µm, respectively). We justified the use of a single kernel given the relatively narrow distribution of best kernels across neurons of each type and the fact that for individual neurons the exact kernel had a modest effect on the calculated dissimilarity of the relevant spike trains. Note that the corresponding analyses based on the best kernels specified for each neuron individually showed virtually the same results.

Across speed discrimination accuracy based on convolved spike trains was similar to that based on firing rate profiles represented in the spatial domain for SA-1 neurons and for FA-1 neurons it was overall better (**Figure 6A**). Mean discrimination accuracy was 48% (range as a function of speed: 34-55%) for FA-1 neurons and 32% (range: 26-36%) for SA-1 neurons. A two-way ANOVA applied to the proportion of correctly classified edge orientations revealed a significant intercept referenced to chance (25%) performance (F_1,51_ = 92.2, p < 0.001, ηp^2^ = 0.64) and main effects of neuron type (F_1,51_ = 20.5, p < 0.001, ηp^2^ = 0.29) and speed (F_7,357_ = 60.3, p < 0.001, ηp^2^ = 0.54). On average across neurons, the discrimination accuracy was generally better for FA-1 than for SA-1 neurons. Regarding the speed effect, for both neuron types, the discrimination accuracy was fairly uniform for speeds up to 45 mm/s, after which it gradually decreased with increasing speed. This decrease was stronger for FA-1 than for SA-1 neurons, which rendered a significant interaction between speed and neuron type (F_7,357_ = 8.7, p < 0.01, ηp^2^ = 0.15).

**Figure 6.**
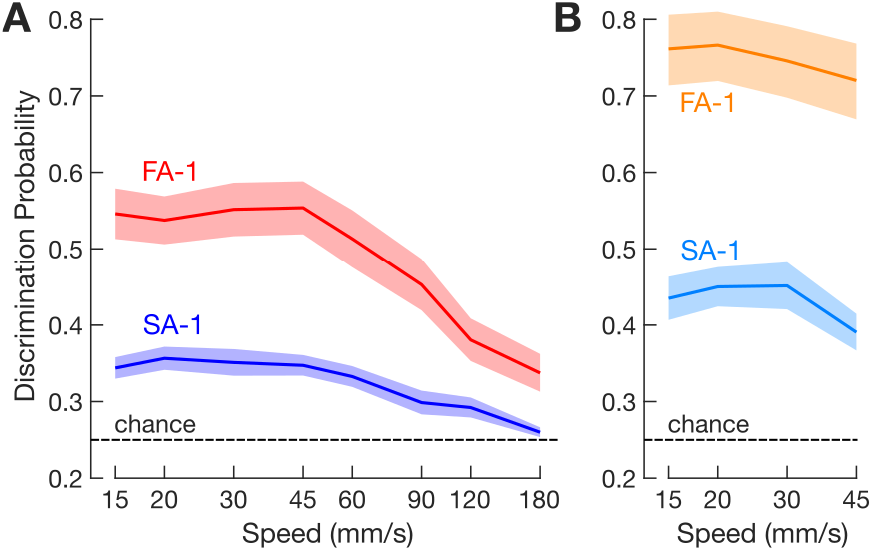
**(A)** Average across-speed discrimination accuracy for FA-1 (red line) and SA-1 neurons (blue line) as a function of scanning speed. (**B**) Same format but considering only the speeds commonly used by people when discriminating these kinds of stimuli (15 - 45 mm/s). (**A** and **B**) Data based on spike trains convolved by the average best kernel over neurons and speeds (66 μm for FA-1 neurons, n = 30; 73 μm for SA-1 neurons, n = 23). Shaded areas illustrate the standard error of the mean and the dashed line indicates chance discrimination performance (25%).

From a behavioral perspective, the above analyses of speed-invariant signaling includes scanning speeds that span a much wider range (15 - 180 mm/s) than what people commonly use (15 - 45 mm/s) when discriminating similar stimuli (Olczak et al., 2018; Vega-Bermudez et al., 1991). Therefore, to provide a more pragmatic view on neurons’ speed-invariant signaling, we calculated across speed discrimination accuracy limited to the range of these commonly used speeds (**Figure 6B**). When considering the commonly used speeds, edge orientation discrimination accuracy was srikingly high, averaging 75% (range as a function of speed: 72-77%) for FA-1 neurons and 43% (range: 39-45%) for SA-1 neurons. A two-way ANOVA focusing on the commonly used speeds revealed a significant main effect of speed (F_3,153_ = 12.5, p < 0.001, ηp^2^ = 0.20) and neuron type (F_1,51_ = 5.6, p < 0.001, ηp^2^ = 0.33) but no significant interaction (F_3,153_ = 0.13, p = 0.72). That is, the superiority of FA-1 neurons was uniform across the commonly used speeds.

Our analyses so far have concerned data pooled over all the edge orientation differences present on our stimulation surface (Δθ = 5°, 10°, 15° and 20°). Discrimination accuracy for all combinations of edge orientations and speeds is illustrated as a confusion matrix in **Figure 7**. To directly address how orientation difference influences the ability of FA-1 and SA-1 neurons to signal edge orientation, we examined how across-speed discrimination accuracy was affected by orientation difference for each pair-wise orientation difference and speed (chance = 0.5). As expected, discrimination accuracy increased gradually with increasing orientation difference (**Figure 8**). A three-way ANOVA with orientation difference and speed as within-group (repeated measures) factors and neuron type as between-group factor indicated a main effect of orientation difference on discrimination accuracy (F_3,153_ = 35.2 p < 0.001, ηp^2^ = 0.41). The superiority of FA-1 neurons over the SA-1 neurons remained (F_1,51_ = 15.0, p < 0.001, ηp^2^ = 0.23) as did the observation that discrimination accuracy varies with speed (F_7,357_ = 21.4, p < 0.001, ηp^2^ = 0.30). This effect appeared consistent across neuron types as we found no significant interaction between scanning speed and neuron type (F_7,1071_ = 1.5, p = 0.15, ηp^2^ = 0.03). We did, however, find a significant interaction between scanning speed and orientation difference (F_21,1071_ = 1.8, p< 0.05, ηp^2^ = 0.03) consistent with the observation that the effect of orientation difference on discrimination accuracy decreased with increasing speed.

**Figure 7.**
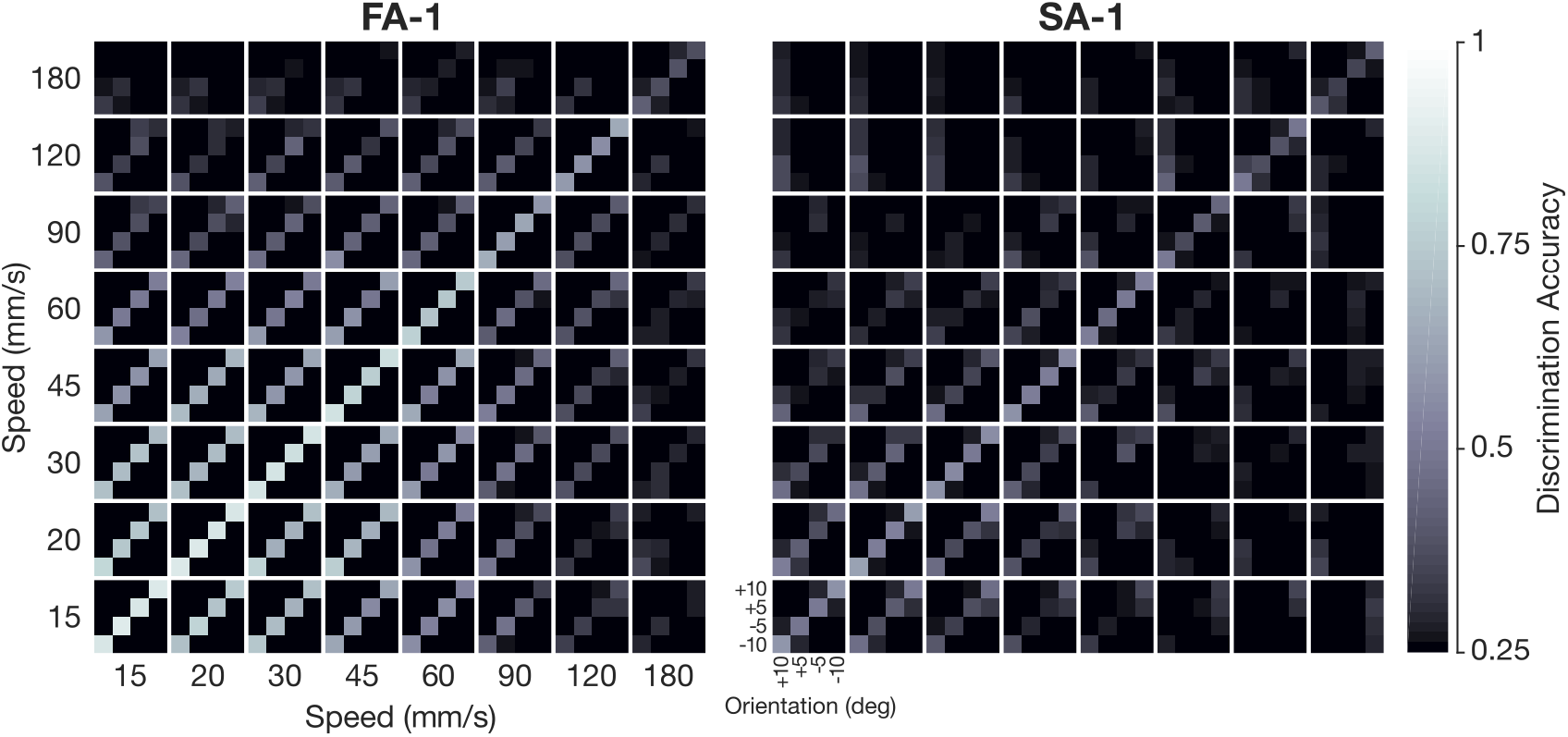
Discrimination accuracy for all edge-orientation and speed combinations based on spike trains convolved by the average best kernel over neurons and speeds (66 μm for FA-1 neurons, n = 30; 73 μm for SA -1, n = 23). Each 4×4 sub-matrix represents comparisons of the four oriented edges within the indicated speed combination. Thus, the main diagonal of the submatrices represents within-speed comparisons, and the off-diagonal submatrices represent across-speed comparisons. The brightness of each element represents the probability of assigning the correct edge-orientation to each of the four possibilities where chance performance is 25%.

**Figure 8.**
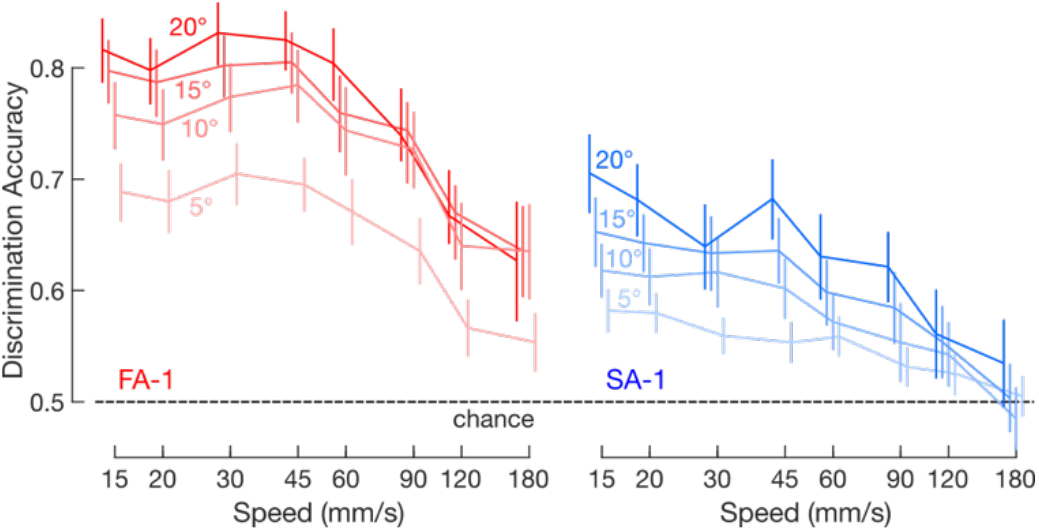
Mean across-speed discrimination accuracy for FA-1 (red lines) and SA-1 neurons (blue lines) as a function of speed and orientation difference. Data based on spike trains convolved by the average best kernel over neurons and speeds (66 μm for FA-1 neurons, n = 30; 73 μm for SA -1, n = 23). The different shades of each color represent the indicated orientation differences. Error bars represent the standard error of the mean and the dashed line indicates chance discrimination performance (50%).

Lastly, given our interest in signaling very fine edge orientation differences, we asked at what scanning speeds individual FA-1 and SA-1 neurons could discriminate the finest orientation difference in our dataset (Δθ = 5°). That is, for each neuron type we examined if discrimination accuracy exceeded chance performance (50%) by using a one-tailed one-sample t-tests for each speed followed by Bonferroni correction for multiple comparison (N = 8). For FA-1 neurons, discrimination accuracy exceeded chance performance for all but the highest two speeds (t_29_ 4.41, p_corrected_ ≤ 0.005 for speeds ≤ 90 mm/s; t_29_ < 2.6, p_corrected_ > 0.06 for speeds >90 mm/s). And for SA-1 neurons, discrimination accuracy exceeded chance performance for all but the three highest speeds (t_22_ ≥ 3.2, p_corrected_ ≤ 0.02 for speeds from ≤ 60 mm/s; t_22_ ≤ 1-8, p_corrected_ ≥ 0.38 for speeds ≥ 60 mm/s). Thus, at scanning speeds that are commonly used in tactile spatial discrimination tasks, individual neurons of both types can contribute speed-invariant information about edge orientation at a level of detail corresponding to differences of only 5°.

## Discussion

We show that individual FA-1 and SA-1 neurons carry substantial information about fine orientation differences in the sequential structure of their spike trains but little in the intensity of their responses. Moreover, the sequential structure allows speed-invariant signaling of fine edge orientation differences across a wide range of scanning speeds when represented in the spatial domain, when action potentials are referenced to the position of the edge stimulus on the skin. In contrast, response intensity is informative only when a neuron is stimulated with a uniform scanning speed.

Our findings reveal that human first-order tactile neurons signal edge orientation most reliably at slow to moderate speeds, corresponding to those used when performing tactile spatial discrimination tasks with the fingertips (Olczak et al., 2018; Vega-Bermudez et al., 1991). We speculate that the choice of speeds in such tasks is associated with the ability of first-order tactile neurons to produce the relevant information. Edge orientation sensitivity at natural speeds was strikingly high, with neurons showing speed-invariant orientation signaling for differences as small as 5°, which is substantially better than what humans can consciously report (Bensmaia et al., 2008; Lechelt, 1992; Olczak et al., 2018; Peters et al., 2015) and on a par with what people exhibit in object manipulation (Pruszynski et al., 2018).

The edge orientation sensitivity we report is likely an underestimate of neurons’ actual sensitivity. First, many first-order tactile neurons could reliably discriminate even our most finely spaced edge orientations. Second, our analysis ignored the viscoelastic and anisotropic properties of the fingertip, and thus possible complex time-varying deformations of the fingertip skin as a surface slides over it (Delhaye et al., 2019; Jarocka et al., 2021). Varying amounts of compression, stretching and shearing of the skin within a neuron’s receptive field may therefore have occurred during the repeated scans both within and across speeds and between edges with different orientations. Such events would have erroneously increased our estimate of the spatial jitter of action potentials across the repetitions of a given stimulus and thus erroneously reduced our estimate of edge orientation information in the spike trains.

For stimuli with larger orientation differences (≥ 22.5°), we previously showed that a neuron’s sensitivity to edge orientation was linked to the layout of subfields in its receptive field (Pruszynski & Johansson, 2014). That is, we could fairly well predict the structure of the generated spike trains (in terms of the presence of spikes and the lengths of gaps between spike bursts) as a function of edge orientation based on how the neuron’s subfields were sequentially stimulated. In the present study we could not directly relate a neuron’s edge orientation signaling to the sensitivity topography of the receptive field because we did not collect receptive field maps. Nevertheless, two pieces of evidence suggest that the mechanism is the same. First, that a neuron exhibits speed-invariance in signaling fine edge orientation differences especially at natural scanning speeds is consistent with its responses being structured by its subfield layout, which for FA-1 and SA-1 neurons is rather speed invariant at least for speeds up to 60 mm/s (Jarocka et al., 2021). Second, the estimate of the spatial precision of action potentials in this study is practically identical to the spatial acuity of the subfield arrangement of the FA-1 and SA-1 neurons as mapped by small dots laterally scanning the receptive field (Jarocka et al., 2021). That is, in both studies, the estimated spatial precision for both neuron types corresponds to the dimension of single fingerprint ridges.

FA-1 neurons were generally better than SA-1 neurons in signaling fine edge orientation based on the spatial structuring of their responses. A plausible explanation is that the receptive fields of human FA-1 neurons have about twice as many subfields as SA-1 neurons (Jarocka et al., 2021; Johansson, 1978; Phillips et al., 1992) while the overall receptive field sizes are comparable (Johansson & Vallbo, 1980). A larger number of subfields could in principle provide higher sensitivity to fine orientation differences by offering more varied sequential structuring with orientation changes (Pruszynski et al., 2018). The superiority of FA-1 neurons in this regard, together with their density in the fingertip being approximately twice as high as that of the SA-1 neurons (Johansson & Vallbo, 1979), suggests that FA-1 neurons play a primary role in extracting detailed spatial properties associated with dynamic fingertip events. Indeed, FA-1 neurons (or those like FA-1 neurons) dominate innervation in other body areas, such as the lips/tongue (Trulsson & Essick, 1997) and foot sole (Corniani & Saal, 2020), where the high-fidelity extraction of dynamic mechanical events seems key to their function. In contrast, the low-threshold mechanoreceptive innervation of the human hairy skin appears dominated by slowly adapting neurons (Corniani & Saal, 2020).

Our study highlights that individual FA-1 and SA-1 neurons can signal fine edge orientation differences via the sequential structure of evoked spike trains. However, for a host of reasons, it seems unlikely that the central nervous system would rely on registering the sequential structure of spike trains in individual neurons to estimate edge orientation. For example, such a scheme would require the spike train to be considered in its entirety which would take a long time especially at slower scanning speeds due to the sizable spread of a neuron’s subfields over the skin and it would not function at all without lateral scanning motion. And for speed invariant signaling of edge orientation information, it would require that the nervous system have knowledge about the time course of each spike train. Similar shortcomings would also apply to estimating edge orientation information based on intensity measures such as peak/mean firing rate and spike counts at the level of individual neurons.

We have instead proposed a population level mechanism for geometric feature signaling that utilizes the high spatial precision of spike generation offered by the presence FA-1 and SA-1 neuron subfields and which can signal tactile spatial details both instantaneously and in a speed invariant manner (Hay & Pruszynski, 2020; Pruszynski et al., 2018; Pruszynski & Johansson, 2014). Briefly, the basis is that subfields belonging to different neurons are highly intermingled because of the high density of FA-1 and SA-1 neurons especially in the fingertip skin (Johansson & Vallbo, 1979) and the substantial size of the skin area in which a neuron’s subfields are distributed (Jarocka et al., 2021; Johansson & Vallbo, 1980). Consequently, when an edge of a certain orientation deforms the skin at a particular location, a subset of neurons whose subfields spatially coincide with the edge are synchronously excited, but at a slightly different orientation or location synchrony occurs primarily in another subset of neurons. In this scheme, the degree to which stimuli synchronously engage different subsets of neurons determines the ultimate edge orientation resolution of the nervous system as well as its resolution in the case of discrimination of tactile spatial details in general. Our previous simulations based on realistic innervation density and receptive field sizes suggest that the presence of subfields is particularly beneficial for edge-orientation signaling for very short edges that involve only a small part of the fingertip which might be particularly relevant of the precise control of the digits in tasks that require dexterous object manipulation (Pruszynski et al., 2018).

The present study cannot ultimately determine how the nervous system builds and propagates information about the high-dimensional space of stimulus features required for doing object manipulation or exploratory tasks. However, the anatomical and functional organization of tactile ascending pathways, as well as sensory pathways in general, suggest that synchronized activity across neuronal ensembles induced by sensory events may play a key role (Brette, 2012; Bruno, 2011; Pruszynski & Zylberberg, 2019; Salinas & Sejnowski, 2001; Singer, 1999; Stanley, 2013). A hallmark of sensory ascending pathways, including tactile pathways, is massive increases in divergence and convergence as excitatory connections synapse sequentially and downstream representations engage orders of magnitude more neurons than the incoming axons (Babadi & Sompolinsky, 2014; Jones, 2000). This dimensionality expansion together with the subfield organization of first-order neurons and the sensitivity of neuronal activity to the timing of synaptic input would offer exceptional potential for simultaneous high-resolution identification of various tactile spatiotemporal events, based on patterns of correlated inputs across neurons. Furthermore, different synaptic integration times imply that both the precise timing of individual spikes in presynaptic axons as well as the intensity of spike bursts could contribute depending on the time scales of the sensory inputs and of the nature and context of the current tasks (Harvey et al., 2013; Hay & Pruszynski, 2020; Lankarany et al., 2019). For object manipulation that requires rapid information about changes in spatial contact states, the nervous system could primarily rely on the precise timing of input across neuronal ensembles, while in perceptual judgment tasks, that usually take place under less time pressure allowing more time for synaptic integration, information in individual neurons’ spike intensities could play a greater role (Hay & Pruszynski, 2020; Johansson & Birznieks, 2004; Pruszynski et al., 2018). It is well established that the central processing of sensory information in general can depend on task and context (Crapse & Sommer, 2008; Engel et al., 2001; Gazzaley & Nobre, 2012; Manita et al., 2015; Schroeder et al., 2010; Zagha et al., 2013). For the somatosensory system, descending motor-related signals at different levels of the sensorimotor system can affect virtually all levels of the tactile processing pathways (Adams et al., 2013; Canedo, 1997; Fanselow & Nicolelis, 1999; Ghez & Pisa, 1972; Lee et al., 2008; Manita et al., 2015; Seki & Fetz, 2012; Zagha et al., 2013). A seemingly effective way to adapt tactile feature computations as a function of behavioral goals would be for descending mechanisms to act on synapses of the tactile ascending pathways to influence the exact pattern of synchronized activity across the neuronal ensembles that are propagated. The plausibility of this assumption, we believe, will be clarified by recent advances in recording neural activity from earliest stages of tactile processing in the spinal cord (Confais et al., 2017) and brainstem of awake and behaving animals (Conner et al., 2021; He et al., 2021; Versteeg et al., 2021).

## Acknowledgements

This work was funded by the Swedish Research Council (Project Grant 22209 to J.A.P.) and the Canadian Institutes of Health Research (Foundation Grant to J.A.P.). J.A.P received salary awards from the Human Frontiers Science Program (Long Term Fellowship) and the Canada Research Chairs program. We thank Ewa Jarocka, Carola Hjältén, Göran Westling, Anders Bäckström, Per Utsi and Etay Hay for their technical support.

